# Mutations Causative of CPEO Differentially Engage Innate Immunity Sensors

**DOI:** 10.64898/2026.05.24.727547

**Authors:** Joshua Okletey, Megan Muench, Chenxiao Yu, Alessandra Maresca, Valerio Carelli, Marco Tigano

## Abstract

Chronic Progressive External Ophthalmoplegia (CPEO) is a primary mitochondrial disorder (PMD) caused by mutations in nuclear genes encoding mitochondrial DNA (mtDNA) maintenance proteins. CPEO is characterized by mtDNA depletion and deletions, and patients primarily present with ocular and muscular features (isolated CPEO). However, additional encephalomyopathy, neurological complications, and Parkinsonism can drive a more severe disease form, CPEO-plus. The evolution from isolated CPEO to CPEO-plus remains poorly understood. Inflammatory and innate immune processes are emerging as strong disease modifiers and may underlie this heterogeneity.

Instability of mitochondrial DNA is a major driver of organellar stress and release of mitochondrial contents into the cytosol. Mutations in several genes involved in mtDNA replication and maintenance have been implicated in triggering the escape of mitochondrial nucleic acids from the mitochondrial matrix. Once exposed to cytosolic innate immune sensors, mtDNA and mitochondrial double-stranded RNA (mt-dsRNA) act as potent immunogens, with more than 10 innate immune sensors capable of recognizing them. Therefore, mtDNA and mt-dsRNA release are likely pathological mechanisms in CPEO, yet the list of CPEO-related genes that can trigger inflammatory processes is far from complete.

Here, we use patient-derived fibroblasts from individuals with CPEO carrying mutations in RNASEH1 and Twinkle, and provide – for the first time – evidence that their mutations drive innate immune activation through the release of different mitochondrial nucleic acids. RNASEH1 mutations lead to the accumulation and subsequent release of mt-dsRNA, while mtDNA remains protected. On the other hand, mutations in Twinkle cause the release of mtDNA without triggering mt-dsRNA production, or leakage. Supporting this notion, the POLRMT inhibitor IMT-1, and the STING inhibitor H-151, reduced interferon stimulated genes expression downstream of RNASEH1 and Twinkle mutations, respectively. Further, when we analyzed a unique compound patient line carrying mutations in both genes simultaneously, we detect both species of nucleic acids in its cytosol, indicating that both pathways can be engaged simultaneously in the same cell. Lastly, we show that cytosolic sensing triggers paracrine signaling to activate bystander microglia – the resident macrophages of the retina and brain – with potential implications to the neurological progression of CPEO.

Overall, our findings reveal a new role for RNASEH1 and Twinkle in driving aberrant innate immunity and paracrine inflammation in CPEO. Our data support a model in which innate immunity is a universal feature of mutations causing mtDNA instability; yet different mutations engage distinct sensing pathways, and in complex scenarios multiple pathways can be triggered at the same time. Given the clinical heterogeneity observed in patients with PMDs, our findings that different signaling pathways are triggered in patient-specific manners might have direct implications for precision medicine approaches aimed at targeting specific innate immunity.

## Introduction

Mitochondria evolved from the endosymbiosis of an obligate aerobe by an ancestral archaeon, giving rise to the first eukaryotic cell(*1*). Once considered little more than the cell’s power plants, mitochondria are now recognized as central hubs of cellular life, with roles in calcium and ROS homeostasis, cell death, ageing, and innate immunity(*2–5*). Mitochondrial dysfunction is therefore associated with a wide spectrum of human diseases(*6*). Throughout evolution, mitochondria retained a compact circular genome of ∼16.6 kbp encoding 37 genes, 13 of which code for subunits of four of the five OXPHOS complexes; the remaining ∼1,100 proteins of the mitochondrial proteome are nuclear-encoded. Mitochondrial function is consequently governed jointly by two genomes, and the two compartments continuously exchange signals to coordinate their activities(*7*, *8*). To replicate and maintain their mtDNA, mitochondria rely on a dedicated replisome comprising. POLG, Twinkle, and mtSSBP1 act as polymerase, helicase and single-stranded DNA-binding protein, respectively, and constitute the minimal mitochondrial replisome(*9*). To complete mtDNA replication faithfully, several other functions are required, including DNA and RNA nucleases (MGME1 and RNASEH1), topoisomerases (TOP3A and Top1mt) and ligases (Ligase 3)(*10*).

Given the essential role of mtDNA, mutations in any replisome gene are either embryonic lethal or contribute to Primary Mitochondrial Disorders (PMDs) – devastating conditions affecting approximately 1 in 4,000 newborns, characterized by extreme clinical heterogeneity, diagnostic complexity, and no available cure. Chronic Progressive External Ophthalmoplegia (CPEO) is a PMD characterized by mtDNA instability in the form of deletions and depletions, and the most common genetic causes of CPEO are nuclear mutations in replisome proteins(*11*). The most common mutated gene is POLG gene, but as many as 18 different mitochondrial proteins have been implicated in CPEO pathology, including the helicase Twinkle and the RNA processing enzyme RNASEH1. The clinical presentation is primarily ocular and muscular (isolated CPEO). However, additional encephalomyopathy, neurological complications, and Parkinsonism can propel a worsened form of the disease, CPEO-plus. What drives progression from CPEO to CPEO-plus remains poorly understood.

A growing body of evidence shows that mtDNA instability can activate innate immune signaling through the cytosolic release of mitochondrial nucleic acids(*12–14*). Both mtDNA and mitochondrial double-stranded RNA (mt-dsRNA) engage cytosolic sensors: mtDNA activates the cGAS–STING pathway(*15*, *16*), while mt-dsRNA – a potent immunogen generated by aberrant mitochondrial transcription or defective RNA processing – trigger Rig-Like-Receptors Signaling pathways that converge on the mitochondrial adaptor MAVS (*17–19*). Collectively, sensing of these mitochondrial DAMPs converges on two main signaling outputs: NF-κB-driven pro-inflammatory gene expression and type I interferon signaling, the latter capable of paracrine activation of bystander cells. Over the last decade, this form of mitochondria-driven innate immune activation has been linked to the pathogenesis of several PMDs(*20*), with systemic or tissue-specific inflammation suggesting that impaired mitochondrial function may broadly promote inflammatory signaling, irrespective of the specific mutation or phenotype. However, the mechanisms by which innate immunity is engaged downstream of a given PMD-causing mutation—and whether different mutations, or patients sharing the same mutation, engage different sensors—remain poorly defined. Understanding these specificities is key both to explaining the clinical variability of these conditions and to identifying targeted therapeutic strategies.

Here, we use patient-derived skin fibroblasts from individuals with isolated CPEO and CPEO-plus, carrying mutations in RNASEH1(*21*), Twinkle(*9*), or both. While Twinkle mutations have previously been connected to innate immune activation, we report for the first time that RNASEH1 mutations are causative of aberrant immunity in a human cell model. By employing additional controls and a unique compound line carrying mutations in both genes, we dissect the rules of cytosolic sensor engagement across different CPEO genotypes. Our findings support a model in which innate immune activation is a general feature of CPEO, but the sensors engaged are mutation-specific—even when the upstream mitochondrial phenotypes are shared.

## Results

### CPEO-Causing Mutations in RNASEH1 and Twinkle Impair Mitochondrial Function in Patient-Derived Fibroblasts

To model the immunological consequences of CPEO-associated mutations in RNASEH1 and TWNK, we cultured four patient-derived skin fibroblast lines (P1–P4) alongside two healthy controls (C1–C2); genotypes, mutations, and zygosity are reported in Fig 1A. P1 and P2 were isolated from two relatives with P1 carrying a single heterozygous RNASEH1*^Y29C^* mutant allele and was clinically unaffected, while P2 carried the same mutant allele RNASEH1*^Y29C^* in homozygosis and presented with CPEO-plus. A third RNASEH1 line (P3) carries a homozygous nonsense mutation at codon 98 (p.Arg98*) together with a heterozygous dominant Twinkle*^R303Q^* mutation. This patient was diagnosed with isolated CPEO. To our knowledge, P3 is the first reported compound line combining RNASEH1*^R98*^* and Twinkle*^R303Q^* mutations studied in the context of innate immunity. P4 carries a dominant Twinkle*^R303W^* variant in heterozygosis and was isolated from a patient with CPEO-plus. This final line allows us to separate the contribution of Twinkle from RNASEH1 in driving innate immune responses. C1 and C2 are fibroblasts from healthy individuals and were employed controls throughout the manuscript.

**Figure 1.**
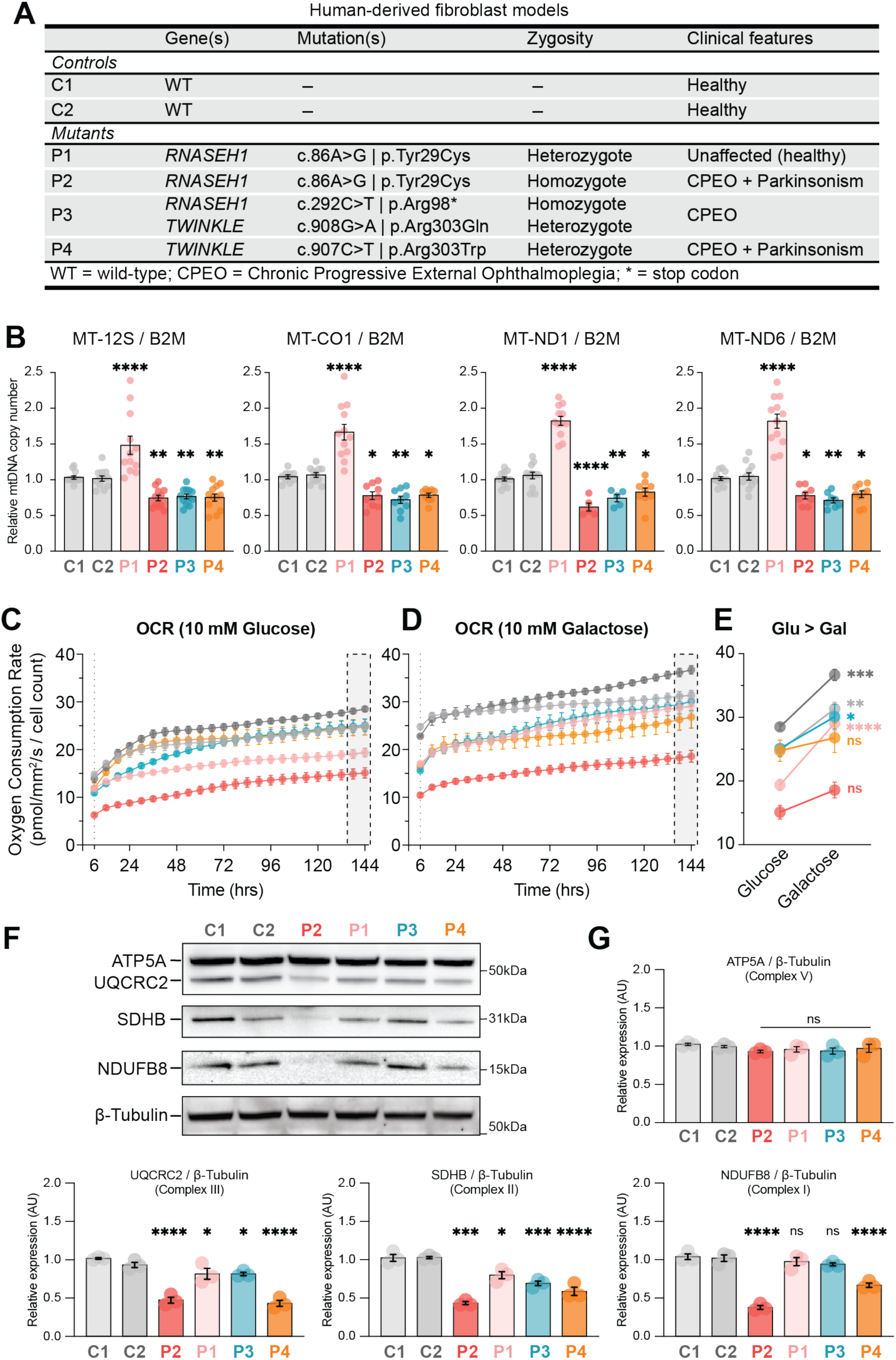
CPEO-associated RNASEH1 and TWNK mutations alter mtDNA copy number and mitochondrial function in patient-derived fibroblasts. (A) Table of human-derived dermal fibroblast models, two control fibroblast cell lines from healthy donors, and four fibroblast cell lines from patients with identified mutations in CPEO-associated genes. Mutation in RNASEH1 were identified in clinically healthy P1 (heterozygous) and P2 (homozygous) diagnosed with CPEO-plus. CPEO-diagnosed P3 harbored mutations in RNASEH1 (homozygous) and TWINKLE (heterozygous), and P4 harbored a heterozygous mutation in TWINKLE. (B) qPCR analysis of mtDNA copy number by relative mtDNA-encoded 12S, MT-CO1/COX I and ND6 to nuclear-encoded B2M. Relative mtDNA copy number are Mean ± SEM derived from n = 4 biological replicates; one-way ANOVA with Dunnett’s post hoc analysis for pairwise comparisons to C1, ∗p ≤ 0.05, ∗∗p < 0.01; ∗∗∗ p≤0.001, ∗∗∗∗p < 0.0001. (C) Resipher measurements of oxygen consumption rate (OCR) in glucose (left) and galactose (center) medium, for 144-hrs, with line plot comparing OCR average of fibroblasts line growing in 96-well plate format and in technical triplicate. Glucose or galactose were added at 10 mM. At the end of experiment, cells were lysed in plate and total protein concentration determined with BCA. The OCR value at the 144 hours was used to generate before-after plots on the right displaying the adaptation to galactose challenge for each cell line. Average of 3 independent experiments is reported and statistical analysis performed with 2-way Anova and Šídák’s post hoc correction. ∗p ≤ 0.05, ∗∗p < 0.01; ∗∗∗ p≤0.001, ∗∗∗∗p < 0.0001. (D) Western blot analysis of nuclear-encoded subunits of the electron transport chain. Equal amounts of lysates were loaded and immunoblotted using the OXPHOS Cocktail antibody (Abcam). (E) Quantification of band intensity, plotted as relative protein expression compared to C1, normalized to β-Tubulin. Mean ± SEM from n -= 3 biological replicates; one-way ANOVA with Dunnett’s post hoc analysis for pairwise comparisons to C1, ∗p ≤ 0.05, ∗∗p < 0.01; ∗∗∗ p≤ 0.001, ∗∗∗∗p < 0.0001.

Published work on patient fibroblasts frequently reports rapid metabolic adaptation, with mitochondrial defects only apparent under additional stress(*22*, *23*). Therefore, we first investigated if mitochondrial dysfunction was detectable across multiple parameters in our patient fibroblasts. Bulk mtDNA copy number was measured by quantitative PCR (RT-qPCR) using primers against three mitochondrial *loci* (MT-12S, MT-CO1,MT-ND1, and MT-ND6), normalized to the nuclear reference gene B2M (Fig 1B). Compared to controls, P1 showed an average ∼1.5-fold increase in mtDNA across all four *loci*, consistent with a compensatory upregulation in mtDNA content. P2, P3, and P4 all showed approximately 30% lower mtDNA content than controls. Mitochondrial transcription mirrored these levels, with ∼30% fewer MT-12S transcripts detected in mutant lines, but not in P1 (Fig S1A). RNASEH1 is essential for the processing of replicative primers from the displacement loop. One of the classical manifestations of its mutation is the increase in an aberrant form of replication intermediate, 7S-RNA. Using a published RT-qPCR assay, we probed the amounts of 7S-RNA in P1, P2 and P3 (all carrying at least 1 mutated allele of RNASEH1) and detected a significant increase in 7S-RNA levels in P2 and P3 (Fig S1B) supporting the pathogenicity of the mutations. P1 displayed no changes, indicating that one intact copy of RNASEH1 is sufficient to process physiological levels of 7S-RNA. Next, to assess respiratory capacity, we monitored oxygen consumption rate (OCR) using the Resipher platform, which record dissolved oxygen in the media of 96-well plates in which live non-permeabilized cells are growing undisturbed(*24*). Recording can be extended over multiple days and normalized to the growth rate of each cell line. In our experiments, 20,000 fibroblasts per line were plated in triplicate in 96-well plates and followed for over six days in glucose-containing media (10 mM), with media changes every 48 hours (Fig 1C). In glucose, P2 showed the lowest OCR among all lines followed by P1, indicating significant respiratory deficiency. On the other hand, P3 and P4 displayed OCR levels that matched those of the controls. To unmask additional deficiencies, we performed Resipher recordings where glucose was substituted with galactose to force oxidative metabolism. In these conditions, all the lines increased their OCR output indicating that galactose is effectively pushing OXPHOS in fibroblasts (Fig 1D). Yet, when we compared the endpoint OCR value between glucose and galactose conditions and normalized these values to total protein content (Fig 1E), P2 and P4 failed to produce a significant increase, while P3 showed a partial but significant compensatory response. P1 increased OCR at a rate higher than that of controls, indicating that one RNASEH1 allele is sufficient to sustain the increased workload imposed by galactose. From Resipher experiments we cannot extrapolate a theoretical ATP production rate (as done for Seahorse). Therefore, to directly quantify ATP production, we used a luminometric assay and measured both total and oligomycin-insensitive (glycolytic) ATP (Fig S1C). Mitochondrial ATP (Total ATP – Oligomycin-insensitive ATP) was significantly reduced in all patient lines, with P2 showing the greatest deficit. Glycolytic ATP was comparable to control levels made exception for P2, where an increase was detected, possibly suggesting a reliance on glycolysis to sustain cell growth. To corroborate OCR and ATP data, we used western blotting and antibodies against representative subunits of complexes I (NDUFB8), II (SDHB), III (UQCRC2), and V (ATP5A)(Fig 1E-F). P2 and P4 showed the strongest decreases in complexes I, II, and III; P3 showed reductions in complexes II and III with a small, non-significant change in complex I; P1 showed only mild changes. None of the patient lines displayed a reduction in the complex V subunit ATP5A.

We then examined the mitochondrial network by immunofluorescence. Fixed cells were stained with anti-TOMM20 antibody and phalloidin to mark the mitochondrial network and cell boundaries, respectively (Fig S1D). Individual cells were segmented and mitochondrial morphology quantified using Mitochondria-Analyzer(*25*). P2 and P4 showed reduced mitochondrial surface area, and signs of mitochondrial fragmentation (lower number of branches and junctions), with P2 displaying the sharpest phenotype (Fig S1E-G). Qualitatively, P2 and P4 also displayed a condensed perinuclear mitochondrial distribution. P3, capable of compensating OCR, also displayed an increase in mass and elongated mitochondria which is indicative of higher performance(*26*). As expected, P1 mitochondria were unchanged.

In summary, four patient-derived fibroblast lines were characterized across a panel of mitochondrial parameters. All symptomatic lines showed approximately 30% reduction in mtDNA content, mirrored by decreased transcription. From a functional perspective, P2 and P4 showed the most consistent impairment while P3 mounted compensatory responses to achieve normal OCR, elevated mitochondrial mass, and an elongated fused network. P1 showed only modest changes across the board, in accordance with the unaffected status.

### CPEO-Related Mutations Broadly Trigger an Inflammatory Response in Human Fibroblasts

To characterize the transcriptional landscape of our CPEO models, we performed unbiased RNA sequencing and differential gene expression analysis. Grouping all CPEO mutants (P2–P4) together, we identified 621 upregulated and 418 downregulated genes relative to the two healthy controls (Fig 2A). Gene Set Enrichment Analysis (GSEA) revealed that five of the top ten enriched pathways were related to inflammatory responses, with TNFA signaling via NF-κB and interferon response pathways being the top two hits (Fig 2B). NF-kB and Type I Interferon responses are the major downstream outputs of cytosolic Damage Associated Molecular Patterns (DAMPs) sensing, which include mitochondrial nucleic acids. We used the log2 fold-changes from a curated set of Interferon Stimulated Genes (ISGs) to create an heatmap, which showed a clear and significant separation between mutant and control lines (Fig 2C). In this case, while P3 was able to compensate the most functions, it displayed the strongest level of ISGs activation. This analysis establishes inflammatory and Type I Interferon responses as a shared transcriptional feature of CPEO. Considering the different background of each line, we also repeated DEG analysis and GSEA stratifying each patient. This analysis revealed a substantial number of both upregulated (ranging from 519 to 1743) and downregulated (ranging from 296 to 1,810) DEGs (Fig S2A-F). Terms related to inflammatory processes remained the most consistently enriched categories, but line-specific responses were also apparent. For example, P2 and P4 showed enrichment of unfolded protein response pathways, and P4 additionally showed glycolysis and apoptosis. Interestingly, P1 showed several transcriptional programs as well, including cholesterol homeostasis, hypoxia, and ROS pathways, suggesting that this mutation triggers broad cellular adaptations even in the absence of overt mitochondrial dysfunction.

**Figure 2.**
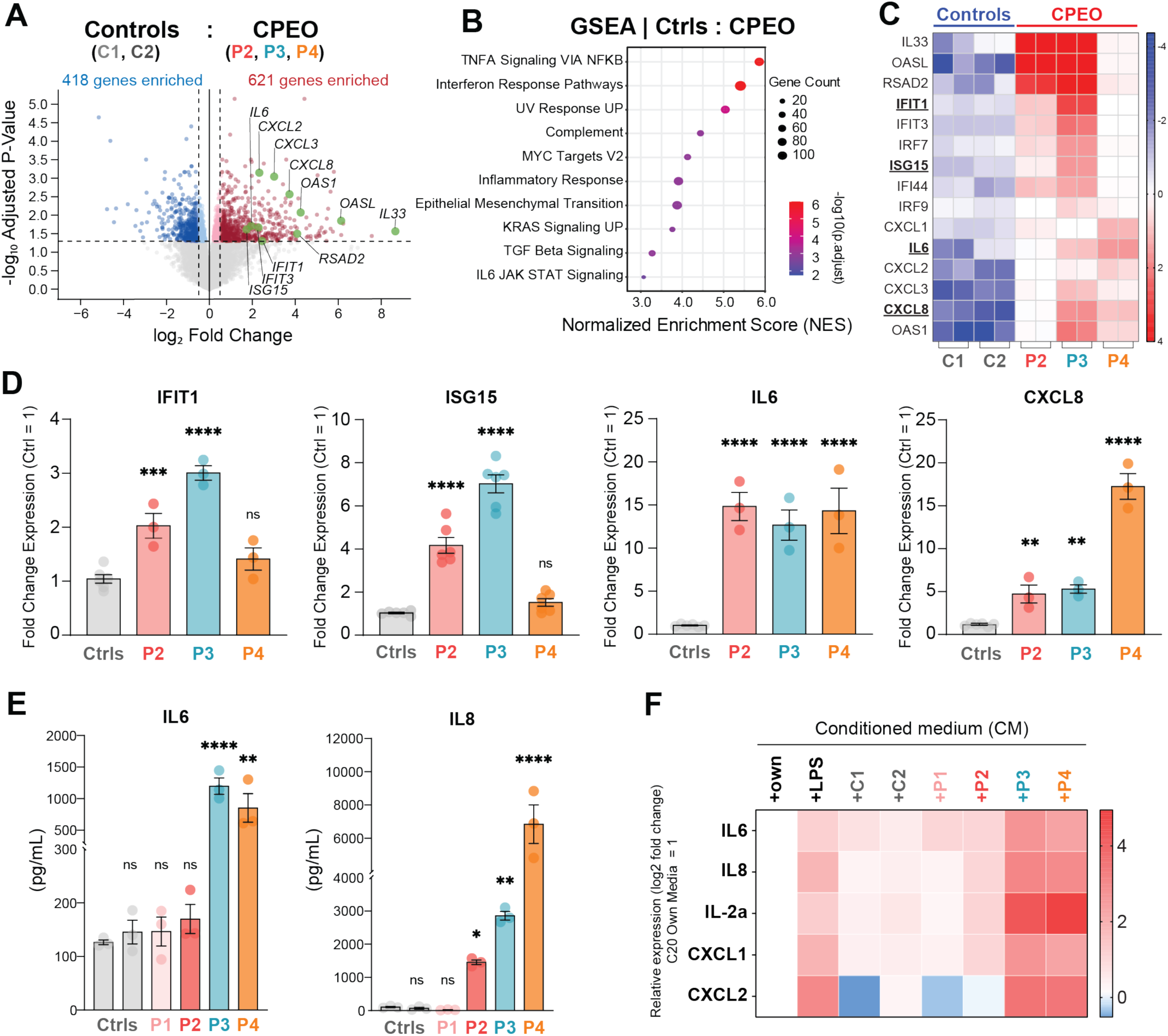
CPEO fibroblasts activate inflammatory and interferon-associated programs and release factors that stimulate microglia. (A) Volcano plot of differential gene expression, comparing CPEO fibroblasts (P2,P3,P4) to Ctrls (C1,C2). Differential expressed genes (DEGs) with fold change ≥ 1.5 with adjusted p-value > 0.05. Highlighting significant DEGs implicated in innate immunity and inflammatory response pathways. (B) Gene Set Enrichment Analysis (GSEA) of genes differentially expressed in CPEO fibroblasts compared with controls, showing enriched Hallmark pathways ranked by normalized enrichment score. Dot size indicates gene count and color indicates adjusted P value. (C) Heatmap of selected interferon-stimulated and inflammatory genes in control and CPEO fibroblast RNA-seq samples. Each column represents one biological replicate, grouped by cell line. Arrowheads mark genes validated by RT-qPCR. (D) RT-qPCR validation of IFIT1, ISG15, IL6, and CXCL8 expression in CPEO fibroblasts relative to pooled healthy controls. Shown is the Mean ± SEM from at least 3 independent experiments. one-way ANOVA with Dunnett’s post hoc analysis for pairwise comparisons to C1, ∗p ≤ 0.05, ∗∗p < 0.01; ∗∗∗ p≤0.001, ∗∗∗∗p < 0.0001. (E) IL-6 and IL-8 concentrations measured in conditioned or exhausted fibroblast culture media by ProQuantum assays. Shown is the Mean ± SEM from at least 3 independent experiments. One-way ANOVA with Dunnett’s post hoc analysis for pairwise comparisons to C1, ∗p ≤ 0.05, ∗∗p < 0.01; ∗∗∗ p≤0.001, ∗∗∗∗p < 0.0001. (F) Heatmap showing inflammatory gene expression measured by RT-qPCR in C20 microglial cells after 48 h treatment with C20 medium, LPS, or conditioned media from control, unaffected carrier, or CPEO fibroblast cultures. Expression values are shown as log2 fold change.

Given the consistent activation of innate immune pathways across all disease models, we validated our transcriptomic results for a panel of ISGs by RT-qPCR (Fig 2D). Although variability between individual genes and lines was observed, RT-qPCR confirmed the RNA-seq results (Fig 2D). Type I interferon responses usually involve the release of cytokines in the culture media, to recall bystander cells. Therefore, we collected exhaust media from fibroblasts and evaluated IL-6 and IL-8 with specific immunoassay (Fig 2E). Consistent with their transcriptional profile, IL-8 was elevated in all mutant lines while IL-6 was increased in P3 and P4 but not in P2. Once again, P1 showed no significant changes. To assess whether the released factors are functionally active and capable of driving paracrine inflammation, we performed conditioned media transfer experiments using C20 microglia (Fig 2F)(*27*). The choice of microglia is relevant to CPEO pathology, being resident immune cells in both the retina and brain tissue. C20 cells were treated for 48 hours with their own media, control or patient fibroblast conditioned media, and LPS as a positive control. As expected, LPS induced significantly all markers tested (IL-6, IL-8, IL-2a, CXCL1, CXCL2). Media from control fibroblasts caused slight but detectable baseline activation, with P1 displaying a similar pattern. Supporting their transcriptional profile, all CPEO lines consistently activated microglia above the baseline with P3 and P4 eliciting a stark response which was higher than the positive control. P2 also consistently activated 4 out of 5 markers, albeit at lower magnitude.

In conclusion, while responses differed between lines, taken together, our transcriptomic and cytokine data confirm that type I interferon responses and NF-κB-driven pro-inflammatory signaling are common outputs of both RNASEH1 and Twinkle mutations. To our knowledge, this is the first time that RNASEH1 and Twinkle mutations have been interrogated in the context of aberrant innate immunity in human fibroblasts model of CPEO and CPEO-plus. Critically, we also provide evidence that the transcriptional profile of the tested mutants translates into release of active cytokines in the media which could transfer inflammatory signals to microglia, a phenomenon that could be relevant to progression from CPEO to CPEO-plus.

### RNASEH1 and Twinkle Mutations Cause the Release of Mitochondrial Nucleic Acids into the Cytosol

Mitochondrial nucleic acids, when in the cytosol, can engage multiple innate immune sensors depending on their backbone, subcellular location, and the cell type. Having established that strong inflammatory responses are common in our CPEO lines, we asked whether mitochondrial nucleic acids are released into the cytosol. RNASEH1 activity is required to degrade RNA:DNA hybrid intermediates of mtDNA replication, and its mutation led to their aberrant accumulation. Therefore, we first tested whether RNASEH1 loss leads to aberrant accumulation of mt-dsRNA. To detect mt-dsRNA we used the specific antibody J2 – capable of detecting double-stranded RNA structures with high specificity(*28*, *29*)] – while nuclei were counterstained with DAPI, and mitochondria with anti-TOMM20 antibodies (Fig 3A). To validate the specificity of J2 in human fibroblasts, we performed a pilot staining in presence and absence of RNASE-III, which should lead to the abrogation of any J2 signal, when specific. This analysis confirmed that the signal identified was of specific nature (Fig S3A). Compared to controls, P2 and P3 showed a striking accumulation of intracellular dsRNA. By contrast, P1 showed no dsRNA, also consistent with the absence of inflammatory responses. P4, carrying an exclusive Twinkle mutation, showed no detectable dsRNA, suggesting that the inflammatory phenotypes driven by Twinkle could be of different nature. Quantification confirmed that P2 and P3 each accumulated on average of ∼40 discrete J2-positive structures per cell. Further, we used TOMM20 and DAPI to calculate the amount of cytosolic dsRNA, which accounted to approximately 40-50% of the total (Fig 3B), and represent the pool of RNA accessible to RNA-sensors, including RIG-I and MDA5(*17*, *30*). Although J2 immunofluorescence is highly specific for dsRNA, it does not distinguish its cellular origin. To confirm that at least a part of the dsRNA detected by J2 in CPEO lines had mitochondrial origin, we treated cells with the POLRMT inhibitor IMT-1 (5 µM, 24 hours), which transiently depletes mitochondrial transcription(*31*). IMT-1 depleted MT-12S transcripts by ∼90% in control cells and ∼50% in P2 and P3 (Fig S3B). Importantly, we also detected a corresponding ∼50% reduction in total J2-positive dsRNA foci in both lines (Fig S3C), suggesting that the detected dsRNA structures are indeed of mitochondrial origin.

**Figure 3.**
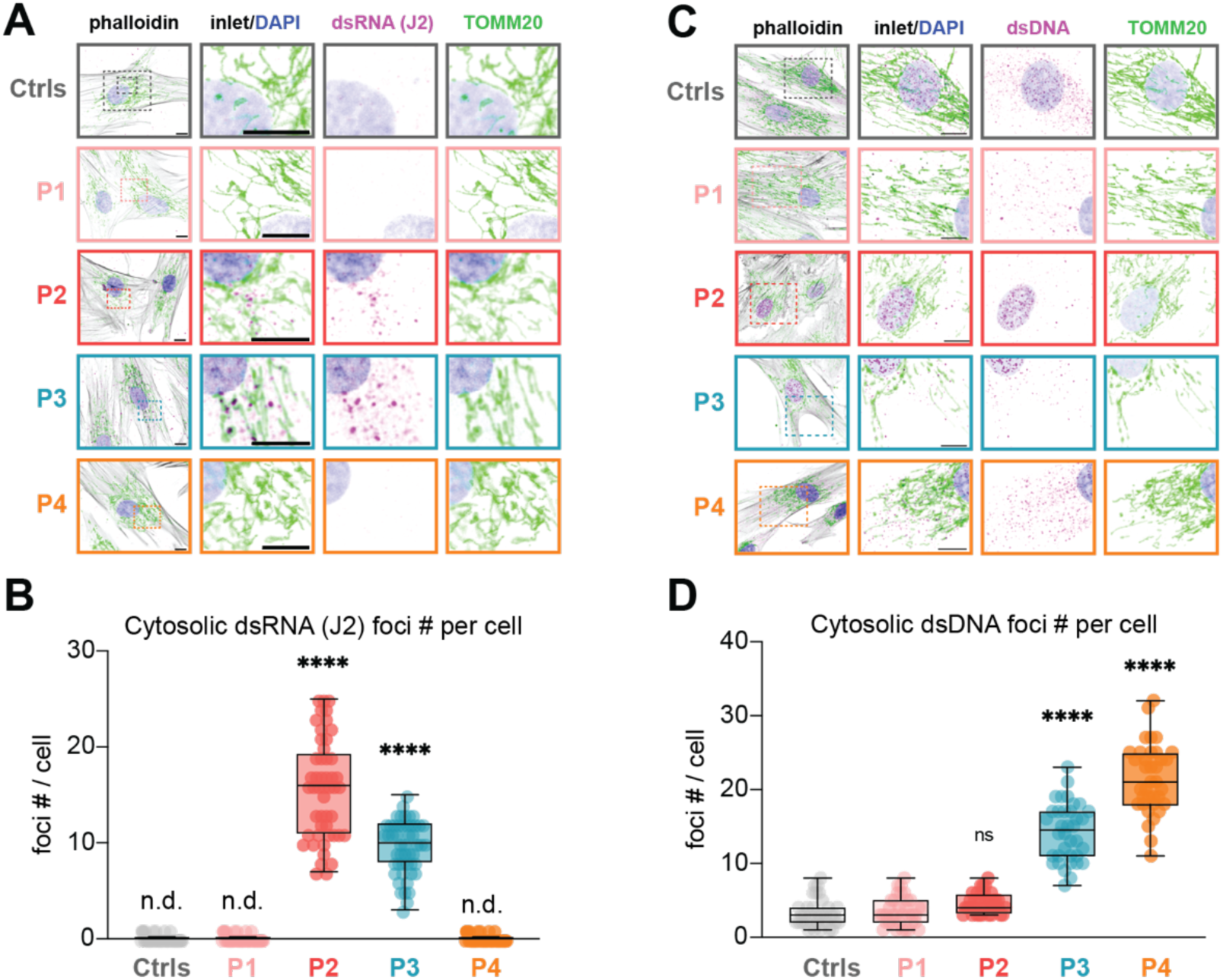
RNASEH1 and TWNK mutant fibroblasts accumulate cytosolic dsRNA and dsDNA. (A) Representative immunofluorescence images of control, unaffected RNASEH1 carrier, and CPEO fibroblasts stained with DAPI (nucleus), phalloidin, J2 antibody to detect dsRNA, and TOMM20 to mark mitochondria. Insets show magnified regions from the boxed areas. Scale bars, X µm; insets, X µm. (B) Quantification of cytosolic J2-positive dsRNA foci per cell in control, P1, P2, P3, and P4 fibroblasts. Cytosolic foci were defined as J2-positive foci outside TOMM20-positive mitochondrial masks and outside DAPI-positive nuclear masks and counted from 50 cells. Ordinary One-way ANOVA with Tukey’s post hoc for pairwise comparisons to C1, ∗p ≤ 0.05, ∗∗p < 0.01; ∗∗∗ p≤0.001, ∗∗∗∗p < 0.0001. (C) Representative immunofluorescence images of control, unaffected RNASEH1 carrier, and CPEO fibroblasts stained with DAPI, phalloidin, anti-dsDNA antibody to detect dsDNA, and TOMM20 to mark mitochondria. Insets show magnified regions from the boxed areas. Scale bars, X µm; insets, X µm. (D) Quantification of cytosolic dsDNA foci per cell in control, P1, P2, P3, and P4 fibroblasts. Cytosolic foci were defined as dsDNA positive foci outside TOMM20-positive mitochondrial masks and outside DAPI-positive nuclear masks and counted from 36 cells. Ordinary One-way ANOVA with Tukey’s post hoc analysis for pairwise comparisons to C1, ∗p ≤ 0.05, ∗∗p < 0.01; ∗∗∗ p≤0.001, ∗∗∗∗p < 0.0001.

While P2 and P3 accumulated cytosolic mt-dsRNA, P4 did not while still mounting a robust inflammatory response. Given the role of Twinkle in unwinding mtDNA during replication and previous reports pointing to a role of Twinkle mutations in the delivery of mtDNA to autophagic compartments(*32*), we hypothesized that mtDNA might gaining access to the cytosol in our fibroblasts. Using a validated anti-dsDNA antibody that detects mitochondrial DNA in immunofluorescence(*33*), we stained all patient lines and quantified dsDNA foci located outside mitochondrial boundaries. P4 showed a consistent accumulation of cytosolic dsDNA (Fig 3E), placing it in location where it can engage sensors such as cGAS. Intriguingly, P3 showed both cytosolic dsRNA and cytosolic dsDNA together with the strongest inflammatory profile. These observations are consistent with a model in which RNASEH1 mutation drives RNA sensing via mt-dsRNA, Twinkle mutation drives DNA sensing via cytosolic mtDNA, and the compound P3 line engages both pathways simultaneously, producing an amplified response.

### CPEO Mutant Fibroblasts Differentially Engage RNA and DNA Sensors

Having established that cytosolic mt-dsRNA and dsDNA are present downstream of RNASEH1 and Twinkle, respectively, we asked whether blocking RNA- or DNA-sensing could rescue innate immune activation. To test the role of RNA sensing downstream of RNASEH1 mutations, we decided to deplete the trigger at the source by blocking mt-RNA transcription with IMT-1. P2 and P3 fibroblasts were treated with IMT-1 (5 µM, 24 hours), after which total RNA was collected and ISG expression assessed by RT-qPCR. ISG expression was substantially reduced in both P2 and P3 (Fig 4A), consistent with mt-dsRNA being the primary driver of immune activation downstream of RNASEH1 mutation. To block DNA sensing, we used the STING inhibitor H-151, which blocks the responses downstream of cGAS. H-151 treatment did not reduce ISG expression in P2 but did so in P3 and P4 (Fig 4B), confirming that cGAS–STING-dependent DNA sensing drives the response in Twinkle mutant cells. Together, these inhibitor experiments are consistent with the data obtained with immunofluorescence, and indicate that mt-RNA and dsDNA engage with two distinct arms of the innate immune pathways.

**Figure 4.**
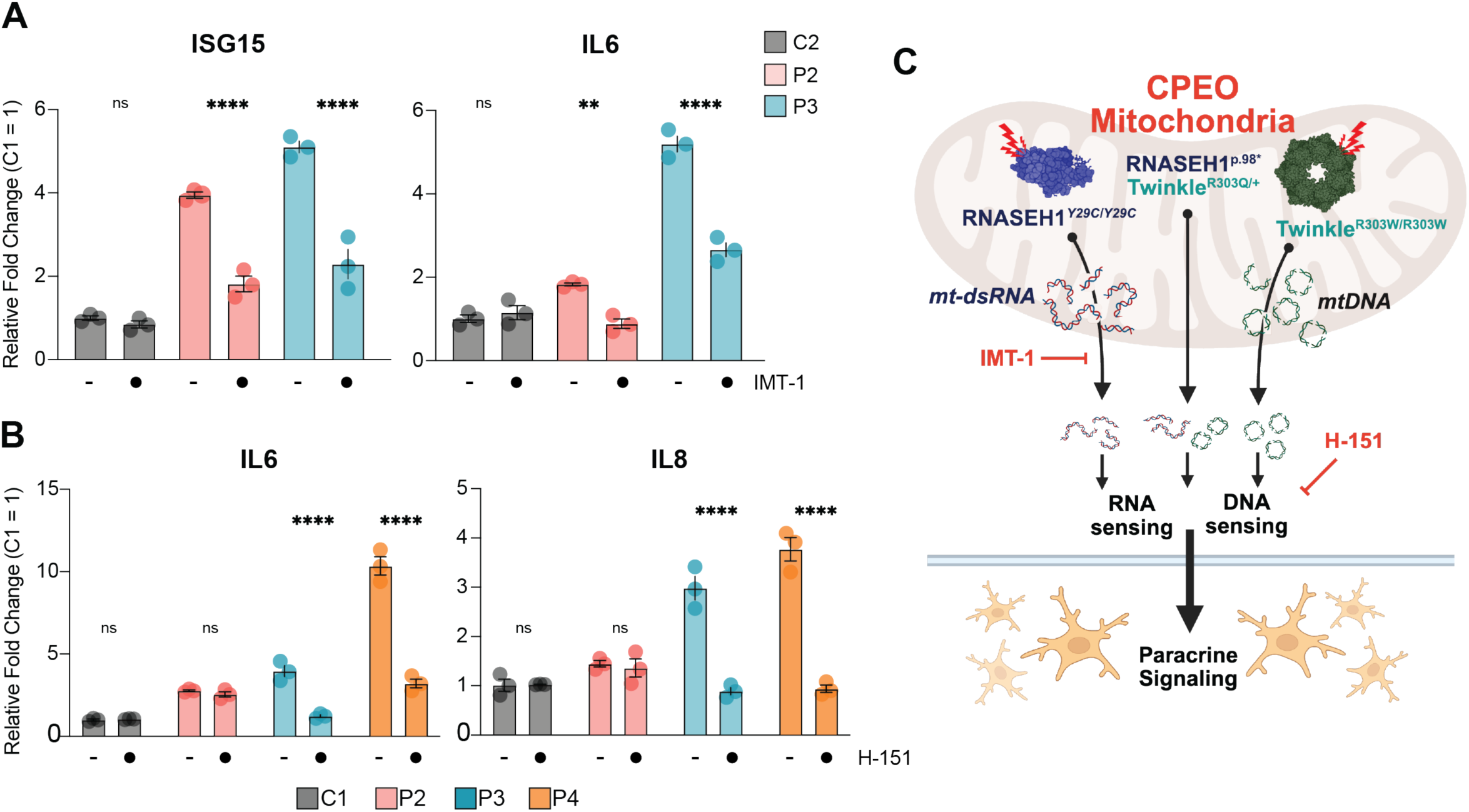
IMT-1 reduces cytosolic dsRNA and inflammatory gene expression in CPEO fibroblasts. (A) Quantification of average J2-positive dsRNA foci per cell from 50 cells after IMT-1 treatment (5 µM, 24 h) in control, P2, and P3 fibroblasts, or no treatment. Ordinary One-way ANOVA with Tukey’s post hoc analysis for pairwise comparisons to C1, ∗p ≤ 0.05, ∗∗p < 0.01; ∗∗∗ p≤0.001, ∗∗∗∗p < 0.0001. (A) RT-qPCR analysis of ISG15 and IL6 expression after untreated or IMT-1 treatment (5 µM, 24 h) in control, P2, and P3 fibroblasts. IMT-1 reduced ISG15 and IL6 expression in P2 and P3 cells. Shown is the Mean ± SEM from 3 independent experiments. 2-way ANOVA with Šídák correction for multiple comparisons. ∗p ≤ 0.05, ∗∗p < 0.01; ∗∗∗ p≤0.001, ∗∗∗∗p < 0.0001. (B) RT-qPCR analysis of IL6 and IL8 expression after untreated or IMT-1 treatment (5 uM, 24 h) in control, P2, P3, and P4 fibroblasts. Shown is the Mean ± SEM from 3 independent experiments. 2-way ANOVA with Šídák correction for multiple comparisons. ∗p ≤ 0.05, ∗∗p < 0.01; ∗∗∗ p≤0.001, ∗∗∗∗p < 0.0001. (C) Graphical Model of differential engagement of innate immunity sensors downstream of RNASEH1 and Twinkle mutations. Created in BioRender. Tigano, M. (2026). Crystal structures of Twinkle and RNASEH1 structures were downloaded from PDB (7T8B; 6VRD). https://BioRender.com/0inufos.

In conclusion, strong inflammatory signatures were detected in all CPEO lines that also had mitochondrial dysfunction phenotypes. Both mt-dsRNA and mtDNA were identified in the cytosol of CPEO and triggered the RNA- and DNA-sensors. In lines with a single mutation, we found only one type of nucleic acid, and the selective inhibition of the relevant sensor was sufficient to rescue ISG expression. In a compound harboring two different mutations, both dsRNA and mtDNA were found in the cytosol, with an amplified response that will likely require a combinatorial inhibition of both sensing pathways.

## Discussion

We characterized four patient-derived skin fibroblast lines harboring mutations in RNASEH1 and Twinkle, two genes implicated in CPEO pathogenesis. RNASEH1 and Twinkle have well-characterized roles in mtDNA replication but – until now – have not been formally tested for a causative role in triggering aberrant innate immunity in human cells. Our set of patient fibroblasts was controlled by two healthy donor lines, allowing us to dissect the contribution of each gene individually and, through a unique compound line, in combination. We also had access to a line from an unaffected relative carrying a single mutated RNASEH1 allele, which allowed us to assess the effects of haploinsufficiency.

Patient fibroblasts are a powerful tool for investigating the role of pathogenic mutations in innate immune activation, though this model comes with caveats. Standardized collection and culturing cannot be guaranteed across different centers and laboratories, and this variability can play an important role when inconsistent results are reported, especially when studying inflammatory outputs(*34*). In our experience, passage number from line establishment is an essential factor influencing cellular adaptation, particularly in inflammatory responses — yet the exact passage number is not always available or reported. All experiments in this manuscript were performed within a maximum passage limit of 12. Beyond adaptation, fibroblasts cannot fully recapitulate the tissue-specific consequences of each mutation. They can be reprogrammed to induced pluripotent stem cells (iPSCs) and used to derive disease-relevant lineages — retinal ganglion cells, cortical or dopaminergic neurons, skeletal muscle — but reprogramming itself may be influenced by the mtDNA status of the donor fibroblast, and may in turn reshape the organellar DNA mutational landscape(*35*, *36*).

Under standardized culturing conditions, we confirmed mitochondrial phenotypes consistent with published data(*21*, *37*). Respiratory profiling also revealed that different mutant lines adapt through distinct — and currently unexplored — mechanisms in response to deleterious mutations(*38*, *39*). For example, the homozygous RNASEH1^Y29C^ line had severely impaired OCR that could not be further increased by galactose challenge. By contrast, the Twinkle^R303Q^ line had OCR comparable to wild-type but similarly failed to increase it with galactose challenge. This is consistent with OXPHOS operating at or near maximum capacity, or in a decoupled mode — a hypothesis that will need high-resolution respirometry to resolve. The decoupling of electrons from ATP synthase activity could also partly explain the reduced levels of ATP production in the same lines. Interestingly, the compound RNASEH1/Twinkle line displayed a respiratory phenotype closer to that of the Twinkle-only line, suggesting that helicase mutations trigger compensatory responses not present in other genetic backgrounds.

The contribution of innate immunity and inflammatory biomarkers to the etiology of primary mitochondrial disorders is being investigated clinically(*40*), propelled by more than 15 years of foundational work connecting mitochondrial contents to cytosolic innate immune sensors. Multiple groups have provided experimental evidence that specific protein structures(*41*, *42*), metabolites(*43*), and nucleic acids(*15*, *17*) – to cite a few – are recognized as DAMPs when they reach the cytosol. While these pathways are now being confirmed in larger human cohort studies (*44*) the number of nuclear-encoded genes formally tested for their contribution to mitochondrial innate immunity remains surprisingly short. Among the nuclear genes with direct ties to mtDNA stability, only a few have been formally tested in patient-derived or animal models. To our knowledge these includes POLG, MGME1, OPA1, TOP3A, TFAM, and TOP1mt (*13*, *15*, *20*, *44–46*). Therefore, we are the first to report that pathogenic mutations in RNASEH1 and Twinkle drive innate immune activation in human patient-derived cells. We acknowledge that Twinkle function has been investigated for its role in promoting the delivery of organellar DNA to endosomes, but not in terms of its outcome on cytosolic sensing of mtDNA(*32*). As for RNASEH1, a connection can be drawn from its nuclear counterpart RNASEH2, which degrades RNA:DNA hybrids. When mutated, RNASEH2 causes Aicardi-Goutières syndrome, characterized by innate immune activation through cGAS-STING (*47*). By analogy, one might expect RNASEH1 mutations to trigger accumulation of mitochondrial RNA:DNA hybrids and activate cGAS-STING — a model that could be supported by the elevated 7S-RNA levels we observed(*48*). Yet, our data obtained with H-151 points to an exclusive role of mt-dsRNA and likely RNA sensing downstream of RNASEH1. At this time, we cannot exclude a previously unrecognized role for RNA:DNA hybrids in triggering RNA sensors or in being recognized by the J2 antibody(*29*, *49*).

Our work adds to the growing list of genetic perturbations driving innate immune activation downstream of mitochondrial dysfunction, complementing other approaches including targeted mtDNA degradation(*14*) or silencing of RNA processing enzymes(*17*). We are therefore comfortable concluding that mtDNA instability can be considered a universal trigger of aberrant innate immunity. Yet, with up to 10 distinct sensors reported to recognize nucleic acids of mitochondrial origin, the downstream inflammatory responses could vary considerably. Patient-to-patient variability, environmental factors including pollution(*50*)or comorbidities such as infection(*44*, *51*), could all play a role in shaping immune responses to mitochondrial dysfunction. The results from our cohort — albeit limited — highlight mutation-specific rules of sensor engagement, even when starting from similar mitochondrial phenotypes. In particular, homozygous RNASEH1*^Y29C^* engaged immunity through cytosolic mt-dsRNA, while dsDNA was detected only within mitochondrial boundaries. Twinkle*^R303Q^*instead caused cytosolic mtDNA release and STING-mediated immune activation. The combination of both RNASEH1*^Y29C^* and Twinkle*^R303Q^*caused escape of both mtDNA and mt-dsRNA, albeit the toxicity of combinatorial inhibition precluded us from formally testing the consequences. While this pattern might seem obvious given the role of RNASEH1 in resolving RNA:DNA hybrids, we believe it warrants closer inspection. First, the closest analogy — RNASEH2 dysfunction in Aicardi-Goutières — would predict cGAS-STING activation via RNA:DNA hybrid accumulation, not RNA sensing. Second, there is currently no evidence for nucleic acids release mechanisms that selectively discriminate between mtDNA and mt-dsRNA; rather, existing mechanisms have mainly been tested with one species or the other and not through direct comparison(*20*, *52*). Therefore, if RNASEH1 mutation is severe enough to allow escape of nucleic acids, it is not immediately clear why mtDNA is not found in the cytosol. We will address this in future experiments, considering differences in patient genetic background, patient-to-patient variation in enzymes that degrade or mask nucleic acids before DAMPs recognition (Liddicoat 2015, Stetson 2008), and the possible existence of regulatory processes that dictate selective nucleic acid leakage.

In conclusion, the marked clinical variability of primary mitochondrial disorders – spanning age of onset, organ involvement, and genotype–phenotype relationships – remains a major barrier to diagnosis, patient stratification, and therapeutic development. Inflammatory and immune processes, now increasingly recognized as an integral part of PMDs and CPEO, are strong disease modifiers(*53–55*) that could contribute to the progression from CPEO to CPEO-plus, currently unexplained. We observed patient-specific engagement of different innate immune sensors downstream of similar levels of mitochondrial dysfunction. Our data suggest that specific rules of engagement for these pathways exist, and that understanding them should be considered a key step toward interpreting patient-to-patient variability in primary mitochondrial diseases.

### Limitations of the study

While well controlled, our cohort is still limited to 4 patient lines and 2 healthy controls. It is challenging to extrapolate general conclusions from a limited number of samples. Ongoing approaches involve expanding the number of cell lines tested and interpret the results in the light of other confounders that cannot be properly addressed yet, including, but not limited to, sex as a biological variable. Our observations lack mechanistic validation of the typology of sensor engaged downstream of each mutation. While inhibitors of POLRMT and STING are widely cited and used, a direct genetic approach removing each cytosolic sensor will be required to complete the signaling landscape caused by RNASEH1 and Twinkle mutations. Similarly, dsDNA and J2 antibodies have specific controls associated, but do not discriminate sequences. A sequencing approach to fully establish causality will be required.

## Acknowledgments

We are grateful to the entire team at University of Bologna and IRCCS Instituto delle Scienze Neurologiche di Bologna for providing us with cells and guidance, and for the technical and intellectual feedback received regularly throughout the project. We thank Dr. Piera Pasinelli, Dr. Davide Trotti, and Dr. Hristelina Ilieva at the Jefferson Weinberg ALS Center for providing us skin fibroblasts from healthy donors.

This work has been supported by NIH-NIGMS R35GM147191. Joshua Okletey salary is supported by NIH-NIGMS T32GM144302. Chenxiao Yu, Marco Tigano salaries are supported NIH-NIGMS R35GM147191. Sidney Tchiong and Megan Muench salaries are supported by institutional funds.

## Declaration of Generative AI and AI-assisted technologies in the writing process

During the preparation of this work, the authors used ChatGPT/Codex, Claude/Claude code and Edison Scientific to improve the readability, grammar, and clarity of the manuscript. The authors take full responsibility for the content of the publication.

## Material and Methods

### Human-derived fibroblast models

Human patient-derived primary skin fibroblasts carrying CPEO-associated mutations in RNASEH1, TWNK, or both were cultured alongside healthy donor primary skin fibroblast controls. Cell lines were a kind gift from Dr. Valerio Carelli and Dr. Alessandra Maresca. The study included two healthy control lines (C1 and C2), one unaffected heterozygous RNASEH1 carrier line (P1; c.86A\>G, p.Tyr29Cys, heterozygous), one symptomatic RNASEH1 mutant line (P2; c.86A\>G, p.Tyr29Cys, homozygous; CPEO plus Parkinsonism), one compound RNASEH1/TWNK mutant line (P3; RNASEH1 c.292C\>T, p.Arg98\*, homozygous; TWNK c.908G\>A, p.Arg303Gln, heterozygous; CPEO), and one TWNK mutant line (P4; c.907C\>T, p.Arg303Trp, heterozygous; CPEO plus Parkinsonism). P1 and P2 were derived from related individuals. Lines were obtained from punch skin biopsy under Institutional Review Board approval. Lines were de-identified and provided to us with only genotype information.

### Cell culture

Fibroblasts were maintained at 37C in a humidified incubator with 5% CO2 and cultured in high-glucose Dulbecco’s modified Eagle medium with GlutaMAX Supplement (DMEM; Gibco, 10566024) supplemented with 10% fetal bovine serum (SeraPrime), 100 U/mL penicillin-streptomycin (Gibco), 0.1 mM MEM non-essential amino acids (Gibco), 1 mM sodium pyruvate (Gibco), and 50 µg/mL uridine (Sigma-Aldrich). Cells were passaged before confluence, and experiments were performed within 7-12 passages for human fibroblasts. C20 human microglial cells used for conditioned-medium transfer experiments were maintained in DMEM/F12 with 10% fetal bovine serum at 37°C and 5% CO2.

### Pharmacological treatments

Mitochondrial transcription was inhibited with the POLRMT inhibitor IMT-1 at 5 µM for 24h (Sigma Aldrich). This treatment condition was used to assess depletion of J2-positive dsRNA foci and to test rescue of interferon-stimulated gene expression. STING-dependent signaling was inhibited with H-151 (MedChemExpress) at 1 µM for 24 hours.

### RNA isolation and RT-qPCR

Total RNA was purified using the NucleoMag RNA kit (Macherey-Nagel) according to the manufacturer’s instructions, which included Genomic DNA digestion. 300-500 nanograms of RNA were reverse-transcribed using the Maxima H Minus First Strand cDNA Synthesis Kit with dsDNase (Thermo Fisher Scientific). Quantitative PCR (qPCR) was subsequently performed for 40 cycles on an Applied Biosystems QuantStudio 5 Real-Time PCR System. Reactions were conducted in triplicates utilizing PowerUp™ SYBR™ Green Master Mix (Thermo Fisher Scientific) in a total volume of 10 μl with standard cycling conditions. Relative gene expression was normalized using ACTB1 as a housekeeping gene, and all calculations were performed using the Quantstudio software. Primer sequences were retrieved from PrimerDB when available or designed at exon junctions retrieved from the Ensembl database (https://uswest.ensembl.org/index.html). 7S RNA was measured using RT-qPCR as described by Manini et al, 2022 (Manini 2022) starting from RNA, rather than DNA. For conditioned-medium transfer experiments, RNA was isolated from C20 cells after treatment and IL-6, IL-8, IL-2A, CXCL1, and CXCL2 transcript levels were measured by RT-qPCR.

### Genomic DNA isolation and mtDNA copy-number analysis

Purified total genomic DNA was used for mtDNA copy number measurement by qPCR and southern blot. Cell pellets harvested were resuspended in a 400 ml PBS buffer containing 0.2% (w/v) SDS, 5 mM EDTA, and 0.2 mg/ml Proteinase K and incubated at 50 C for 6 hours with constant shaking at 1,000 rpm. DNA was precipitated by adding 0.3 M sodium acetate, pH5.2, and 600 µl isopropanol and kept at −20°C for over 2 hours. Precipitated DNA was centrifuged at 20,000 x g at four C for 30 minutes, followed by a 70% ethanol wash. DNA was then resuspended in the TE buffer (10 mM Tris-HCl, pH8.0, 0.1 mM EDTA) and quantified by Nanodrop. To determine the relative mtDNA copy number by qPCR, sequences specific to mtDNA (MT-RNR1/2) and the nuclear B2M locus (Table S1) were independently amplified from 25 ng of total genomic DNA using the same qPCR protocol described above. The Quantstudio software used the primary relative quantification method with mtDNA as the target and B2M as a reference.

### Oxygen-consumption measurements using Resipher

Oxygen consumption rate (OCR) was measured using the Resipher platform in live, non-permeabilized fibroblasts. Twenty thousand fibroblasts were seeded in 100 µL of media, in triplicate wells of 96-well plates and monitored continuously 6 days. Media was changed every 48 hours. OCR was recorded either in base medium containing 10 mM glucose or in no-glucose medium supplemented with 10 mM galactose to force oxidative metabolism. Longitudinal OCR traces were normalized to cell growth obtained in the same conditions. At the end of experiment, total proteins were collected from the well to perform a final quantification normalized by protein concentration per well.

### ATP measurements

20,000 cells were seeded in an opaque-wall 96-well plate, allowed to adhere for 3 hours, after which triplicate wells were treated with 1uM Oligomycin for 1 hour. Following treatment, plate is subject to a CellTiter-Glo® 2.0 Cell Viability Assay, performed according to manufacturer’s protocol with no deviations. Luminescence readings of the assay plate were performed on a plate reader at 1 sec integration per well. Quantification of mitochondrial-generated ATP was performed by subtracting the readings of Oligomycin-treated (non-mitochondrial ATP) wells from the untreated wells (total cellular ATP).

### Cytokine immunoassays

Media conditioned by fibroblasts for at least 48 hours or from confluent plates was collected. Media was briefly centrifuged at 1200 g for 10 minutes. IL-6 and IL-8 concentrations were evaluated with specific ProQuantum Assays (Thermo Fisher Scientific) following recommendations. Concetrations in the supernatant were determined against a standard curve. Reactions were run in triplicate in a QuantStudio5 (Thermo Fisher Scientific).

### Conditioned-medium transfer to C20 microglia

Conditioned media from healthy control, unaffected carrier, and CPEO fibroblast cultures were collected from confluent plates, filtered through a 22 µm filter syringe to eliminate cells, diluted 1:1 with C-20 media and transferred to a 6-well plate containing 1 × 10^6 C-20 cells. C20 cells treated with their own medium as a negative control, or with 1µg/ml LPS (InvivoGen) as positive control. After 48hrs, RNA was collected from C20 cells and IL-6, IL-8, IL-2A, CXCL1, and CXCL2 expression was measured by RT-qPCR.

### Western blotting

Cells were washed with cold PBS and then lysed with SDS lysis buffer (1% SDS, 10 mM TRIS and 1 mM EDTA) with Halt™ Protease and Phosphatase Inhibitor Cocktail (Thermo Scientific). Lysates were sonicated (Sonic Dismembrator Model 500, Fisher Scientific) for 10 cycles of 3 s on/3 s off at 15% amplitude. Lysates were clarified by centrifugation for 30 min at 14,800 rpm, 4 °C and the supernatant was quantified using the enhanced BCA protocol (Thermo Fisher Scientific, Pierce). Lysates were mixed with 4X laemmli loading buffer (SDS: 8.0%, Glycerol: 40.0%, β-mercaptoethanol: 8.0%, Bromophenol Blue: 0.02%, Tris-HCl: 250 mM) and then boiled at 70° C for 10 mins. An equal amount of proteins were loaded on SDS-PAGE gels and then transferred to a nitrocellulose membrane. Membranes were blocked in EveryBlot Blocking Buffer. Incubation with primary antibodies was performed overnight at 4 °C. Membranes were washed and incubated with HRP-conjugated secondary antibodies at 1:5,000 dilution, developed with Clarity ECL (Biorad) and acquired with a ChemiDoc MP apparatus (Biorad) and ImageLab v.5.2. Antibodies against actin were used as loading control dependent on the protein of interest. A full list of antibodies used in the study and relative dilutions is available in Supplementary Table 1.

### Immunofluorescence staining

Fibroblasts were plated on glass coverslips and fixed with pre-warmed 4% paraformaldehyde for 15 min at room temperature. Cells were washed with PBS, permeabilized with 0.1% Triton X-100 in PBS for 5 min, blocked in blocking buffer containing 1% BSA, 0.2% gelatin, and 0.1% Triton X-100 for 30 to 60 min at room temperature, and incubated with primary antibodies for 1h at RT. After PBS washes, cells were incubated with Alexa Fluor-conjugated secondary antibodies, counterstained with DAPI where indicated, and mounted using ProLong Gold Antifade mountant.

Mitochondrial morphology was assessed by TOMM20 immunostaining, with phalloidin staining used to mark cell boundaries. Double-stranded RNA was detected with the J2 antibody, mitochondria were marked with TOMM20, and nuclei were counterstained with DAPI. Double-stranded DNA foci were detected with an anti-dsDNA antibody Images were acquired on a Leica MICA microscope with a 63X N.A 1.2, water immersion lens.

### Image analysis

Images were preprocessed in Leica LAS X acquisition and analysis software, and all analyzed in Fiji/ImageJ. For all image analysis, Phalloidin intensities were used as cell mask, for quantification of cell surface area. TOM20 intensities were used as mitochondria mask, with threshold and morphometric analyses performed using the Mitochondria Analyzer plugin in Fiji. Foci analyses, for both dsRNA and dsDNA foci, were performed on threshold intensities of optical sections using the particle analyzer tool, with size cutoff >0.25um. Quantification was performed on foci within these parameters and within the cell mask, for total foci counts; and foci within the cell mask but not within the mitochondria mask, for cytosolic foci counts.

### RNA sequencing and bioinformatics

Total RNA from fibroblast lines was submitted to Plasmidsaurus for 3’ end-counting RNA-seq on the Illumina platform, with three biological replicates per line. Libraries were prepared by reverse transcription with a poly(dT)VN primer, followed by tagmentation and amplification with unique dual indices; UMIs were used for deduplication. Reads were trimmed with FastP v0.24.0, aligned to the human reference genome using STAR v2.7, and deduplicated with UMICollapse v1.1.0. Gene expression was quantified with featureCounts (subread v2.1.1) using strand-specific counting against 3’ UTR and exon features. Differential expression was assessed with edgePython v0.2.5. Symptomatic CPEO mutant lines (P2-P4) were compared with healthy control lines (C1-C2), and line-specific comparisons were also performed for individual carrier or patient lines relative to healthy controls. Functional enrichment was performed using GSEApy v0.12 with the MSigDB Hallmark gene set. Interferon-stimulated gene heatmaps were generated from log2 fold-change values for a curated ISG set. RNA-seq differential-expression and enrichment analyses used multiple-testing correction, and adjusted P values or false-discovery rates were reported where applicable.

### Statistical analysis

Statistical analyses were performed using GraphPad Prism 11 unless otherwise indicated. For RT-qPCR experiments, each biological replicate, sample, or gene was analyzed in technical triplicate. For imaging experiments, individual cells are displayed to show cell-to-cell variability, but statistical tests were performed on biological replicate-level summary values when biological replicates were available. Two-group comparisons were analyzed using two-tailed Welch’s t tests unless otherwise indicated. Multi-group comparisons were analyzed using one or 2-way ANOVA with Dunnet’s and Šídák corrections where applicable, and as indicated in figure legends. RNA-seq analyses used adjusted P values or FDR-corrected statistics calculated by Plasmidsaurus as specificied above. Statistical significance was defined as P \< 0.05 unless otherwise stated. Error bars and replicate numbers are defined in each figure legend.

---

**Figure S1.**
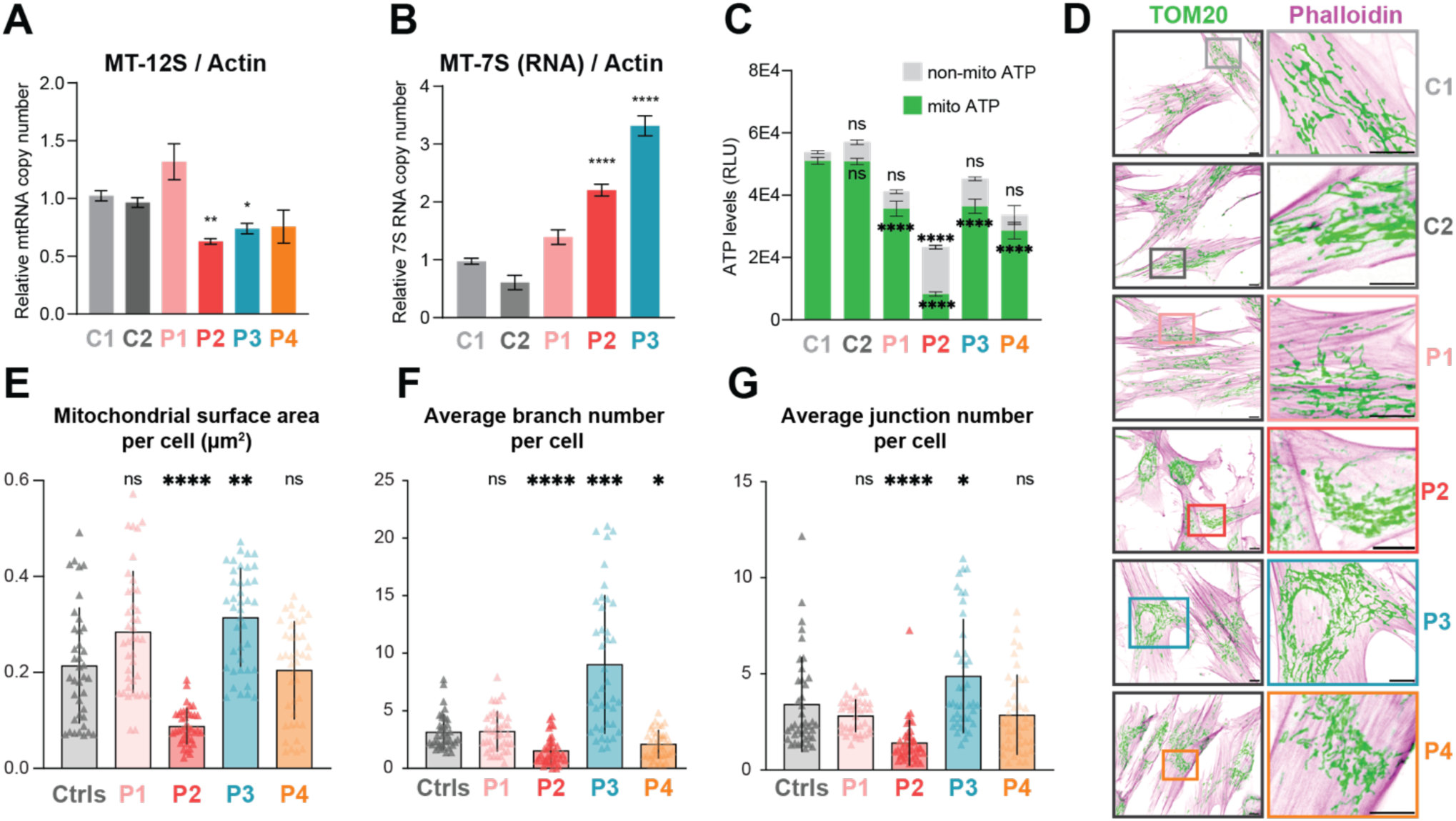
Supporting analysis of mitochondrial transcription, ATP production, and mitochondrial morphology in CPEO fibroblasts. (A) RT-qPCR analysis of MT-12s as a metric of mitochondrial transcription levels in CPEO fibroblasts relative to pooled healthy controls. Shown is the Mean ± SEM from at least 3 independent experiments. One-way ANOVA with Dunnett’s post hoc analysis for pairwise comparisons to C1, ∗p ≤ 0.05, ∗∗p < 0.01; ∗∗∗ p≤0.001, ∗∗∗∗p < 0.0001. (A) Relative abundance of 7S-RNA in indicated CPEO fibroblasts relative to pooled healthy controls. Shown is the Mean ± SEM from at least 3 independent experiments. one-way ANOVA with Dunnett’s post hoc analysis. ∗p ≤ 0.05, ∗∗p < 0.01; ∗∗∗ p≤0.001, ∗∗∗∗p < 0.0001 (C) Total ATP, non-mitochondrial ATP, and mitochondrial ATP measured using a luminescent ATP assay. Non-mitochondrial ATP was defined as oligomycin-insensitive ATP, and mitochondrial ATP was calculated as total ATP minus oligomycin-insensitive ATP. Shown is the Mean ± SEM from; 2-way Anova and Šídák’s post hoc correction. ∗p ≤ 0.05, ∗∗p < 0.01; ∗∗∗ p ≤0.001, ∗∗∗∗p < 0.0001 (D) Representative immunofluorescence images of fibroblasts stained for TOMM20 to mark mitochondria and phalloidin to mark cell boundaries. Insets show mitochondrial network morphology. Scale bar X µm, inset scale bar X µm. (E) Quantification of mitochondrial surface area per cell, average mitochondrial branch number per cell, and average mitochondrial branch junctions per cell from TOMM20/phalloidin images using Mitochondria-Analyzer. Parameters were extrapolated from at least 38-40 cells per cell type and their distribution tested for Normality and Lognormality and statistical significance was calculated with One-way ANOVA Kruskal-Wallis. ∗p ≤ 0.05, ∗∗p < 0.01; ∗∗∗ p≤0.001, ∗∗∗∗p < 0.0001

**Figure S2.**
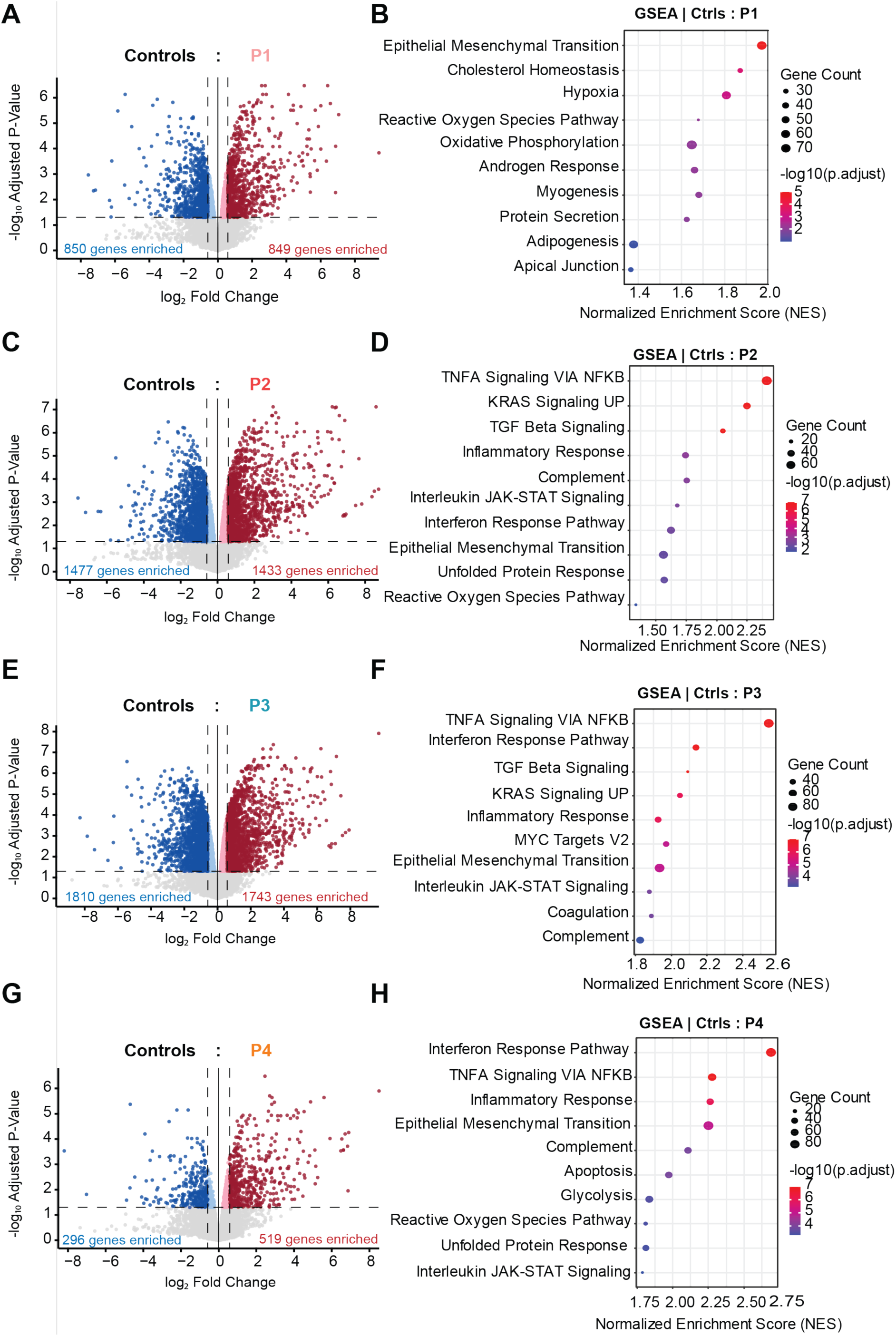
Transcriptomics analysis in control, unaffected carrier, and CPEO fibroblasts. (A, C, E, G) Volcano plots showing differential gene expression for each fibroblast line relative to healthy controls: unaffected RNASEH1 carrier P1 (A), RNASEH1-mutant CPEO plus Parkinsonism line P2 (C), compound RNASEH1/TWNK-mutant CPEO line P3 (E), and TWNK-mutant CPEO plus Parkinsonism line P4 (G). (B, D, F, H) GSEA of differentially expressed genes relative to healthy controls, showing the top enriched MSigDB Hallmark pathways for each comparison. Dot size indicates gene count, and color indicates adjusted P value.

**Figure S3.**
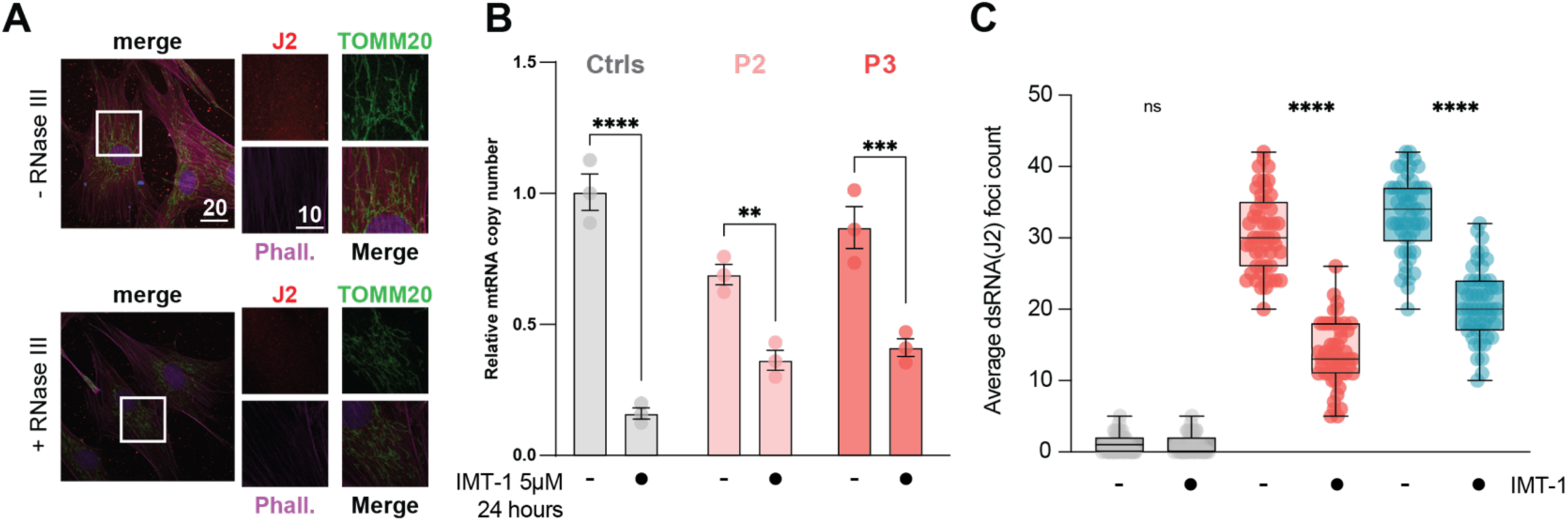
J2 Antibody Staining Specificity and Efficacy of IMT-1 Treatment. (A) Quality control for J2 Specificity. Fibroblasts have higher background dsRNA than other cell lines. To ensure that signal was detected with high specificity, coverslips with fixed P3 cell line were treated with RNASEA before immunostaining with J2. Signal is reverted to that of background, indicating specific binding of dsRNA by the antibody. (B) RT-qPCR analysis of MT-12s normalized to ACT1B gene to evaluate functionality of IMT-1 inhibitor. Cells were treated for 24 hours with 5µM IMT-1 and total RNA was purified and MT-12S relative abundance before and after treatments calculated (Ctrls NT = 1). Shown is the Mean ± SEM from 3 independent experiments. 2-way ANOVA with Šídák post hoc correction. ∗p ≤ 0.05, ∗∗p < 0.01; ∗∗∗ p≤0.001, ∗∗∗∗p < 0.0001. (C) Quantification of average J2-positive dsRNA foci per cell from 50 cells after IMT-1 treatment (5 µM, 24 h) in control, P2, and P3 fibroblasts, or no treatment. Ordinary One-way ANOVA with Tukey’s post hoc analysis for pairwise comparisons to C1, ∗p ≤ 0.05, ∗∗p < 0.01; ∗∗∗ p≤0.001, ∗∗∗∗p < 0.0001.

